# Synaptojanin1 regulates lysosomal functions in ventral midbrain neurons

**DOI:** 10.1101/2022.10.14.512269

**Authors:** Xinyu Zhu, Sanjana Surya Prakash, Geoffrey McAuliffe, Ping-Yue Pan

**Author notes:** **Author contributions:** P-Y. P and X.Z designed the project. P-Y. P supervised the research. S.S.P contributed to plasmid construction. G.M performed the EM experiments. X.Z performed rest of the experiments, analyzed the data, and wrote the paper. X. Z and P-Y. P edited the paper. **Competing Interests:** None.

## Abstract

A major pathological hallmark of Parkinson’s Disease (PD) is the manifestation of Lewy bodies comprised of alpha-synuclein (α-syn). The accumulation of α-syn enriched protein aggregates is thought to arise from dysfunction in degradation systems within the brain. Recently, missense mutations of *SYNJ1* encoding the SAC1 and 5’-phosphatase domains have been found in families with hereditary early-onset Parkinsonism. Previous studies showed that *Synj1* haploinsufficiency (*Synj1+/-*) leads to PD-like behavioral and pathological changes in mice, including the accumulation of the autophagy substrate p62 and pathological α-syn proteins in the midbrain (MB) and striatum. In this study, we aim to investigate the neuronal degradation pathway using the *Synj1+/-* MB culture as a model. Our data suggests that autophagy flux and cumulative autophagosome formation is unaltered at baseline in *Synj1+/-* MB neurons. However, lysosome number is reduced with a similar decrease in lysosomal proteins, including LAMP1, LAMP2, and LAMP2A. Lysosomes are hyperacidified with enhanced enzymatic activity in *Synj1+/-* MB neurons. Using a combination of light and electron microscopy, we show that lysosomal changes are primarily associated with a lack of SAC1 activity. Consistently, expressing the SYNJ1 R258Q mutant in N2a cells reduces the lysosome number. Interestingly, the lysosomal defects in Synj1+/- neurons does not impact the clearance of exogenously expressed wild-type α-syn; however, the clearance of α-syn A53T was impaired in the axons of *Synj1+/-* MB neurons. Taken together, our results suggest axonal vulnerability to lysosomal defects in Synj1 deficient MB neurons.

**Significance Statement:** In the study, Zhu et al. discovered a previously uncharacterized role of Synj1 in regulating lysosomal number, protein, and acidity in ventral midbrain neurons. These alterations are associated with a specific impairment in the clearance of α-syn A53T, but not WT α-syn in axons, suggesting an essential role of Synj1 in axonal degradative capacity under pathological stress. This work in cultured mammalian neurons complements recent research efforts in *Drosophila, C. elegans and* zebra fish, and provides a novel insight for the role Synj1 in neuronal autolysosomal function.

## Introduction

Parkinson’s disease (PD) is characterized by progressive loss of substantia nigra pars compacta (SNpc) dopamine (DA) neurons and the accumulation of Lewy bodies enriched in pathological forms of α-syn (Henderson et al., 2022; Oliveira et al., 2021; Surmeier, Obeso, & Halliday, 2017). While the exact cause of SNpc neuron loss remains elusive, bioinformatics analyses on familial inherited PD and sporadic PD have identified PD genes and risk loci that congregate in the autolysosomal pathway (Fleming et al., 2022; Robak et al., 2017; Udayar, Chen, Sidransky, & Jagasia, 2022; Vazquez-Velez & Zoghbi, 2021). Autophagy (also known as, macroautophagy) is a “self-eating” process that degrades a wide range of cytosolic components including misfolded proteins and damaged organelles to maintain cellular homeostasis (Klionsky, 2007; Nixon, 2013). This process involves the formation of mature autophagosomes followed by fusion with lysosomes for degradation of cytosolic contents. Chaperone-mediated autophagy (CMA) mediated by LAMP2A-assisted protein translocation (Kaushik & Cuervo, 2018; Kon & Cuervo, 2010) is less common but a highly important degradational pathway for α-syn (Cuervo, Stefanis, Fredenburg, Lansbury, & Sulzer, 2004). Whether and how each individual PD gene is implicated in the autophagosome/lysosome dysfunction during pathogenesis remains to be elucidated.

Recently, missense mutations including R258Q, R459P and R839C in PARK20/*SYNJ1* have been discovered in families with early onset parkinsonism (Kirola, Behari, Shishir, & Thelma, 2016; Krebs et al., 2013; Quadri et al., 2013; Taghavi et al., 2018). Reduced *SYNJ1* transcripts were also reported in a subset of sporadic PD brains (Pan et al., 2020). *SYNJ1* encodes synaptojanin1/Synj1, a phosphoinositol phosphatase with two enzymatic domains: a 5’-phosphotase domain that specifically hydrolyzes the 5’-phosphate from the inositol ring of PI(4,5)P_2_ (Guo, Stolz, Lemrow, & York, 1999; Pirruccello & De Camilli, 2012) and a SAC1-like domain that dephosphorylates PI3P, PI4P and PI(3,5)P_2_ into PI (Zhong et al., 2012). Interestingly, Synj1 was initially characterized to cooperate with endophilin to regulate synaptic vesicle recycling through regulating plasma membrane PI(4,5)P_2_ levels and nascent clathrin coated vesicles (Cremona et al., 1999; McPherson et al., 1996; Verstreken et al., 2002; Verstreken et al., 2003). Since the identification of *SYNJ1* mutations in PD, new investigations have emerged that examined the roles of Synj1 in the autolysosomal pathway. Recent studies in *Drosophila and C. elegans* showed that the SAC1 domain is required for autophagosome formation. The R258Q mutation, which abolishes SAC1 enzymatic activity, delays autophagosome maturation due to an accumulation of PI(3)P and PI(3,5)P_2_ and/or disrupted Atg9 activity (Vanhauwaert et al., 2017; Yang et al., 2022). In cultured mouse cortical astrocytes, however, an increase in autophagosome formation was observed in cells from the Synj1 deficient background (Pan et al., 2021). Autophagy impairment was found in the later stages of autolysosomal degradation, and both phosphatase domains were involved in stress-induced clearance of p62 (Pan et al., 2021). Consistently, in aged Synj1 haploinsufficient (Synj1+/-) mice an accumulation of the autophagy substrate p62 was observed in all brain regions including DA neurons in the midbrain (Pan et al., 2020). Additionally, in the cone photoreceptors of zebrafish retina, Synj1 deletion led to mislocalization and abnormal accumulation of late endosomes and autophagosomes (George et al., 2014; George, Hayden, Stanton, & Brockerhoff, 2016). While these recent studies present evidence for a novel role of Synj1 in regulating the autophagic and autolysosomal functions, results obtained from different model systems are somewhat conflicting and the exact step where Synj1 is involved in the autolysosomal degradational pathway in mammalian neurons remains elusive.

*SYNJ1* downregulation was identified in a subset of sporadic PD patients (Pan et al., 2020) and we therefore decided to investigate Synj1 heterozygous (Synj1+/-) deficiency, which represents a more prevalent genetic condition than homozygous point mutations found in few families so far. In this study, we performed a comprehensive analysis of the autophagosome-lysosome pathway in cultured Synj1+/- midbrain neurons. We found that Synj1 haploinsufficiency did not affect the basal level LC3-mediated autophagy flux in the soma of ventral MB neurons. Robust changes were observed in the lysosomal system, including decreased lysosome number, lysosomal proteins, and lysosome hyperacidification. Further, we found the lysosomal defect is associated with a reduction in SAC1 activity rather than 5’-phosphase activity. More interestingly, the degradation of α-syn A53T, but not WT α-syn, was significantly impaired in the axons of Synj1+/- neurons. Taken together, our results indicate specific vulnerability in axonal degradation in response to Synj1 deficiency associated with lysosomal dysfunction.

## Results

### Synj1+/- MB neurons exhibit normal autophagy flux and cumulative autophagosome formation at the basal level

Prior research has shown that in the brains of aged (12-month) male Synj1+/- mice, p62 is increased in the cortex, striatum, and MB DA neurons; however, enhanced LC3B level was only observed in the cortex and striatum but not in the MB DA neurons (Pan et al., 2020). To further investigate whether Synj1 regulates autophagy in MB neurons, we first transfected GFP-LC3 in cultured MB neurons from littermate Synj1+/+ and +/- mice. All analyses were performed in mature neurons between DIV (days in vitro) 13 and 16. Bafilomycin (Baf, 20 nM) was applied to the culture for 1 hour to reveal the basal level autophagy flux. In both MB cultures treated with vehicle or baf, we found similar numbers of GFP-LC3 puncta in the cell bodies between Synj1+/+ and Synj1+/- MB neurons (Fig. 1A-B), suggesting normal basal autophagy flux. GFP-LC3 puncta in the axons could not be determined with confidence in either Synj1+/+ or Synj1+/- neurons and were therefore not included in the analysis. We next employed another molecular probe, mKeima, a pH-sensitive ratiometric fluorescent protein that is resistant to the lysosomal acidic environments and therefore capable of reporting cumulative autophagosome formation and acidification (Katayama, Kogure, Mizushima, Yoshimori, & Miyawaki, 2011). Less acidic (pH6∼pH7) autophagosomes exhibit strong emission at 440 nm excitation but weak emission at 561 nm excitation, while more acidic autophagosomes or autolysosomes (pH4∼pH6) exhibit reverted emission ratio. Due to the lack of quenching, the number of autophagosomes observed at the cell body using mKeima (Fig. 1C-D) represents cumulative autophagosome formation at baseline, which was increased by ∼5-fold compared to those shown by GFP-LC3 (Fig. 1A-B). No significant increase in cumulative autophagosome formation was observed in the Synj1+/- neuronal soma overtime (within 7 days from transfection to imaging) compared to the Synj1+/+ neurons (Fig. 1D). Similarly, mKeima was able to reveal a few more punctate structures along the axons (Fig. 1G) compared to GFP-LC3, however, overall, these puncta were extremely sparse to justify an accurate analysis for axonal density. Thus, we only compared the acidity of autophagosomes/autolysosomes by calculating the ratio of emissions at 440 nm and 561 nm excitations (Fig. 1E-H). We found that Synj1 deficiency did not change the acidity of autophagosomes/autolysosomes either at the soma (Fig. 1F) or the axons (Fig. G-H) of MB neurons. We conclude that in cultured MB neuron, Synj1 does not contribute significantly to autophagosome formation and maturation at the basal level.

**Figure 1.**
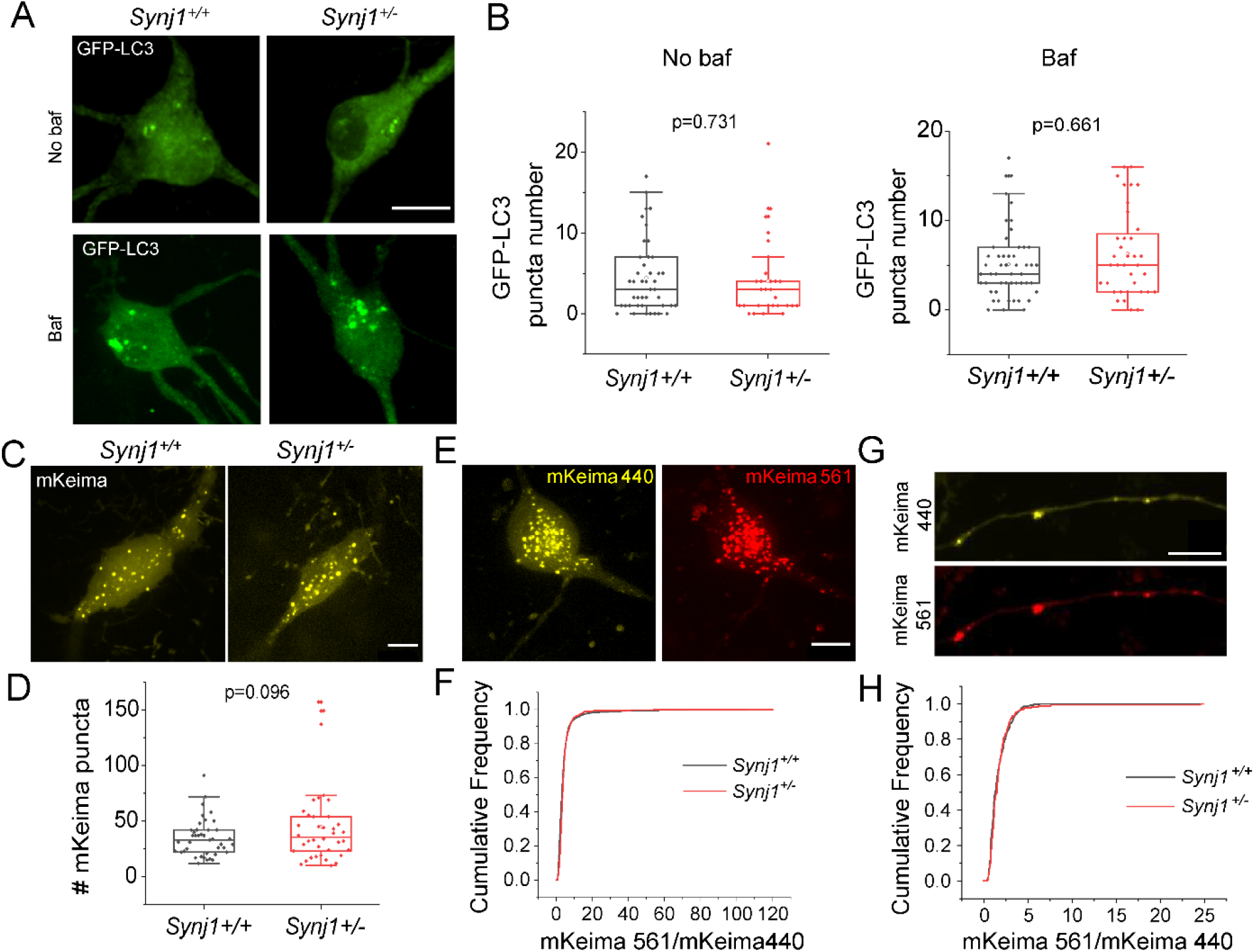
Synj1 deficiency does not substantially alter baseline autophagosome formation and flux in cultured MB neurons. (A)-(B) GFP-LC3 labeled autophagosome in the soma of Synj1+/+ and Synj1+/- MB neurons treated with/without 20 nM bafilomycin A1 for 1 h. (A), representative images. (B), quantification for the GFP-LC3 numbers at the soma. For no baf group, N=44/29 (*Synj1+/+*/*Synj1+/-*) neurons, for baf group, N=51/36 (*Synj1+/+*/*Synj1+/-*) neurons. (C)-(D) mKeima labeled autophagosome in the soma of Synj1+/+ and Synj1+/- MB neurons. (C), representative images. (D), quantification for the mKeima puncta numbers at the soma, N=41/38 (*Synj1+/+*/*Synj1+/-*) neurons. (E) Representative images show mKeima signal in the soma of MB neuron following activation by 440 nM and 561 nM light. (F) Distribution analysis of the ratio (mKeima561/ mKeima440) for the mKeima puncta between Synj1+/+ and Synj1+/- MB neurons, N=2355/899 (*Synj1+/+*/*Synj1+/-*) puncta. (G) Representative images show mKeima signal in the axon of MB neuron from activation by 440 nM and 561 nM. (H) Distribution analysis of the ratio (mKeima561/ mKeima440) for the mKeima puncta between Synj1+/+ and Synj1+/- MB axons, N=297/177 (*Synj1+/+*/*Synj1+/-*) puncta. The *p* values are from student’s *t*-test. Scale bar in all images, 10 μM.

### Synj1 deficiency is associated with changes in the lysosomal system in cultured MB neurons

We next investigated the lysosomal system by examining endogenous expression of lysosome-associated proteins, including LAMP1, LAMP2 and LAMP2A. We found that in the soma of Synj1+/- MB neurons the expression of LAMP1 was reduced by ∼ 25% (Fig. 2A-B). Additionally, the density of LAMP1 structures was significantly lower (Fig. 2C), accompanying a reduction in the immunofluorescence of each LAMP1 puncta in Synj1+/- MB neurons (Fig. 2D), suggesting a possible reduction of lysosome number and protein expression. Recent studies suggested that LAMP1 is not a reliable marker for lysosomes due to its presence on late endosomes (Cheng et al., 2018a, 2018b). We thus performed immunofluorescence for two additional LAMP proteins: LAMP2 and LAMP2A. LAMP2A is a specific isoform of LAMP2 that regulates chaperone mediated autophagy (CMA) (Cuervo & Dice, 1996; Cuervo et al., 2004). Consistently, we found a near 25% reduction in both LAMP2 (Extended Data Fig. 2-1A-B) and LAMP2A (Extended Data Fig. 2-1C-D) at the cell bodies of Synj1+/- MB neurons. In both TH+ and TH-neurons from the midbrain, similar reductions of lysosomal associated proteins were observed (Extended Data Fig. 2-2), suggesting Synj1 deficiency impairs the lysosomal system in ventral MB neurons. Due to the enriched expression of Synj1 in the axons and synapses, we next sought to examine if Synj1 deficiency affects axonal lysosomes. Using immunofluorescence, it was technically challenging to analyze the lysosomes at the axons in the mixed neuron-glia culture. We therefore expressed LAMP1-GFP in cultured MB neurons. Similarly, we found that the LAMP1 puncta density in the axons were significantly reduced in Synj1+/- MB neurons (Fig. 2E-F). Together, these results suggest a previously unknown role of Synj1 in lysosomal biogenesis or maintenance.

**Figure 2.**
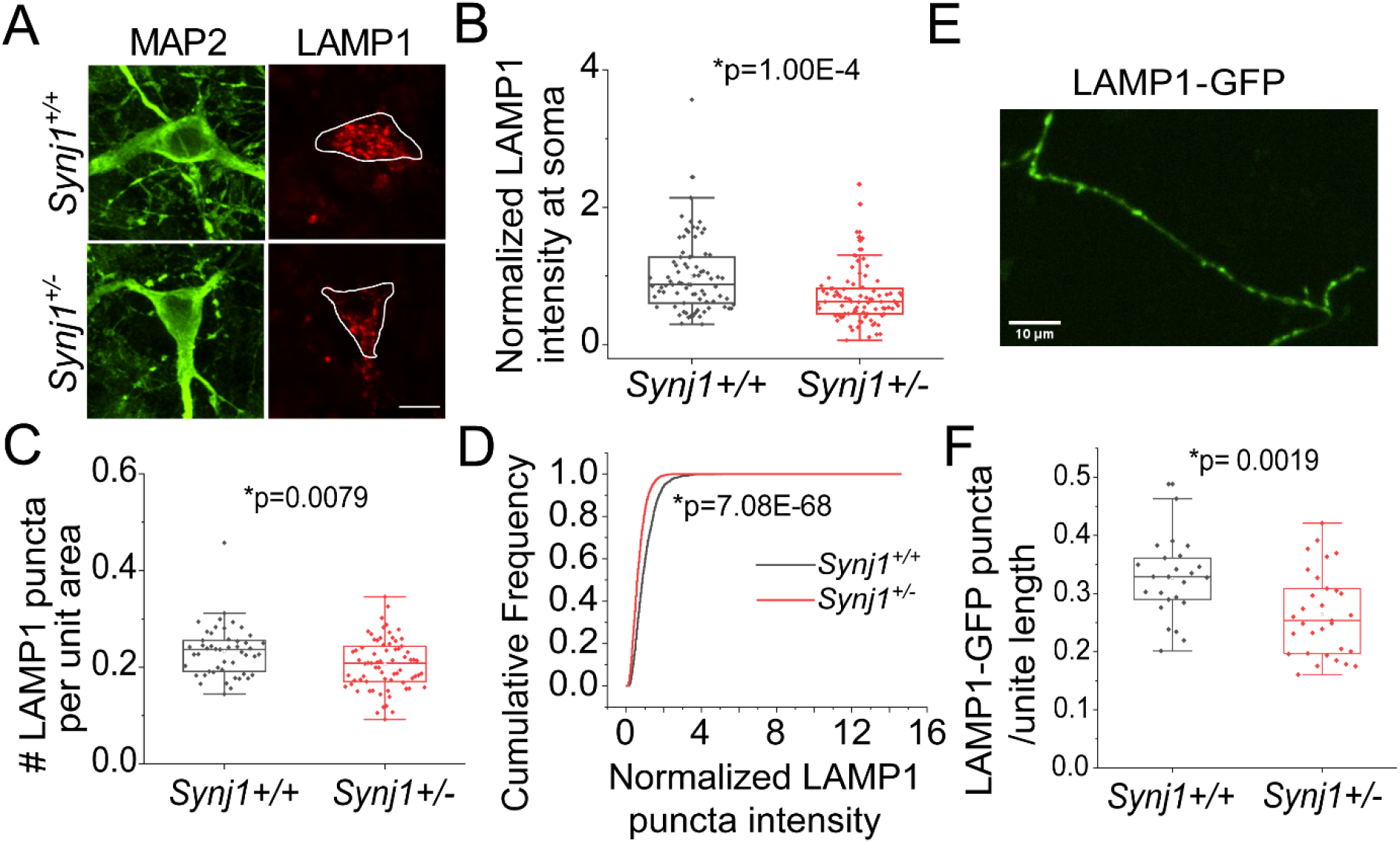
Reduced LAMP1 expression and puncta number in Synj1 deficient MB neurons. (A)-(B) Immunofluorescence analysis of LAMP1 in the soma of Synj1+/+ and Synj1+/- MB neurons, MAP2 is used as neuronal marker. (A), representative images. (B), quantification results, N=82/88 (*Synj1+/+*/*Synj1+/-*) neurons. (C) Quantification of LAMP1 labeled puncta for Synj1+/+ and Synj1+/- MB neurons. Y axis represents the normalized puncta number per unit cytoplasmic area, N=82/88 (*Synj1+/+*/*Synj1+/-*) neurons. (D) Cumulative frequency graph shows the LAMP1 fluorescence intensity distribution of the puncta in soma between Synj1+/+ and Synj1+/- MB neuron. N=2091/2873 (*Synj1+/+*/*Synj1+/-*) puncta. (E)-(F) Analysis of LAMP1-GFP puncta in the axons of Synj1+/+ and Synj1+/- MB neurons. (E), representative images. (F), quantification results, N=25/31 (*Synj1+/+*/*Synj1+/-*) axons. The *p* values for (B), (C) and (F) are from student’s *t*-test. The *p* value for (D) is from Two Sample Kolmogorov-Smirnov test. Scale bar in all images, 10 μM.

### Hyperacidification of lysosomes in Synj1 deficient MB neurons

We next sought to determine whether lysosome function, such as acidification and enzymatic activity, is impaired in Synj1 deficient MB neurons. To investigate lysosomal acidity, we expressed a ratiometric lysosomal pH biosensor, Fluorescence Indicator REporting pH in Lysosomes (FIRE-pHLy, hereafter called pHLy) (Chin et al., 2021). Surprisingly, we found that lysosomal acidity was increased in Synj1 deficient neurons both at the soma (Fig. 3A-B) and at the axon (Fig. 3C-D). The hyperacidified lysosomes may result in elevated enzymatic activity in Synj1 deficient neurons. To test this, we used the Magic Red cathepsin-B assay (MR). Magic red is a substrate of the lysosomal cathepsin-B enzyme that fluoresces red upon cleavage by active cathepsin enzymes. As expected, we found that the MR fluorescence was indeed increased in Synj1+/- neurons (Fig. 3E-F), indicating enhanced lysosomal enzymatic activity.

**Figure 3.**
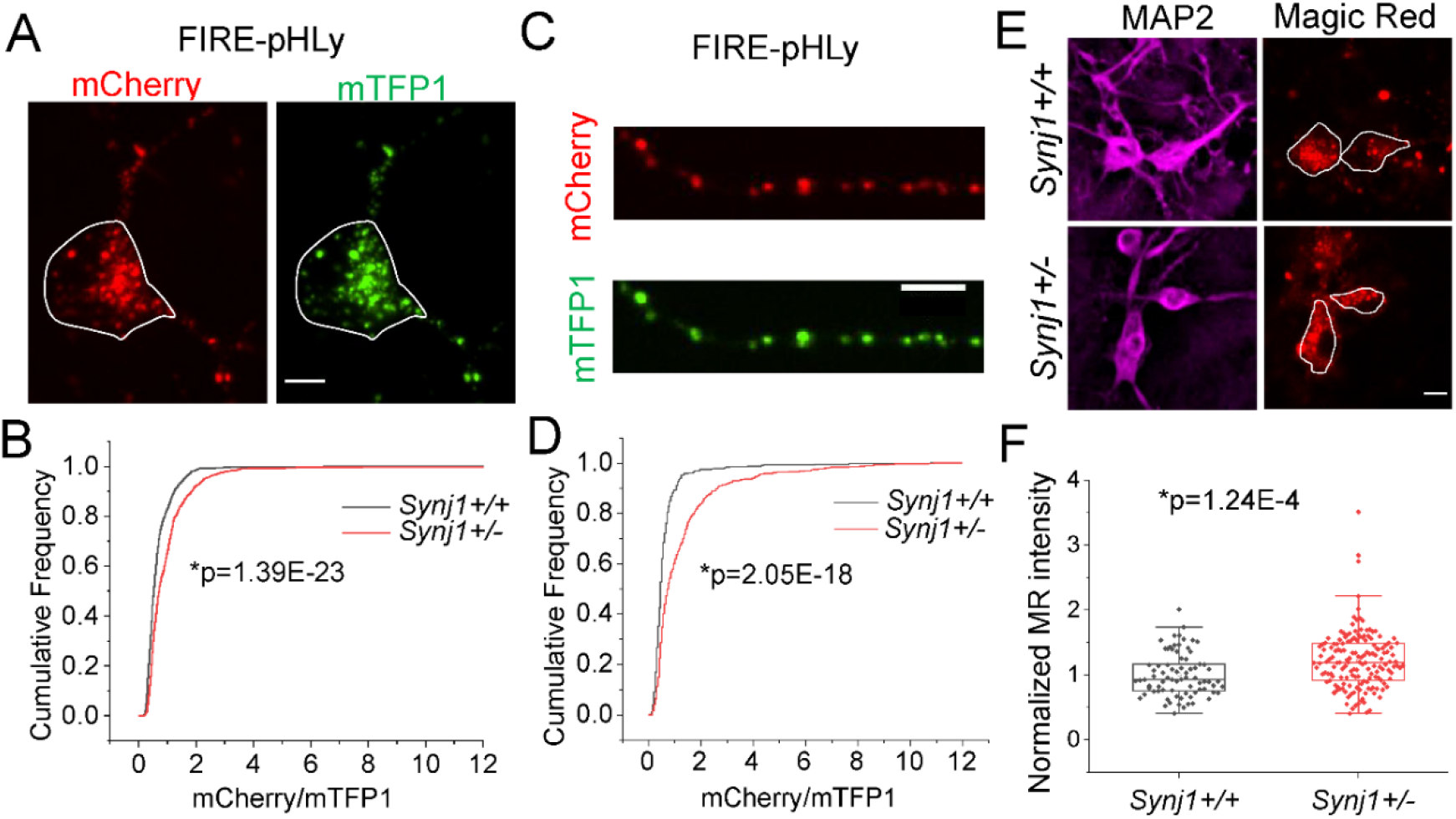
Synj1 deficiency leads to lysosomal hyperacidification and enhanced enzymatic activity. (A)-(B) Lysosome acidity assay using a ratiometric fluorescent probe pHLy. (A), representative images show the mCherry signal and mTFP1 signal in the soma of MB neuron transfected with pHLy. (B), distribution analysis of the ratio (mCherry/mTFP1) for the pHLy labeled puncta between Synj1+/+ and Synj1+/- MB neurons. N=1151/1108 (*Synj1+/+*/*Synj1+/-*) puncta. (C)-(D) Lysosome acidity assay in the axon of MB neuron. (C), representative images show the mCherry signal and mTFP1 signal in axon. (D), distribution analysis of the ratio (mCherry/mTFP1) for the pHLy labeled puncta between Synj1+/+ and Synj1+/- axons. N=432/579 (*Synj1+/+*/*Synj1+/-*) puncta. (E)-(F) Cathepsin-B activity analysis using the dye Magic Red. (E), representative images show the Magic Red signal from live cell imaging and the MAP2 signal from *post hoc* staining for Synj1+/+ and Synj1+/- MB neurons. (F), quantification of the intensity of Magic Red signal between Synj1+/+ and Synj1+/- MB neurons. N=78/159 (*Synj1+/+*/*Synj1+/-*) neurons. The *P* values for (B) and (D) are from Two Sample Kolmogorov-Smirnov test. The *p* value for (F) is from student’s *t*-test. Scale bar in (A) and (C), 5 μM; Scale bar in (E), 10 μM.

### Lysosome defect is associated with Synj1’s SAC1 activity

To further determine a direct association between Synj1 deficiency and lysosomal integrity, and to understand how the two enzymatic domains of Synj1 regulates lysosomes, we expressed wild-type (WT) and mutant human SYNJ1 (SJ1) cDNAs in Neuro 2a (N2a) cells. Among all mutants we examined, SJ1 R258Q and SJ1 R839C were previously identified in Parkinsonism patients (Krebs et al., 2013; Quadri et al., 2013; Taghavi et al., 2018). The R258Q mutation abolishes SAC1 activity, and the R839C mutation reduces both SAC1 and 5’-phosphatase activities (Pan et al., 2020). The SJ1 D769A is not disease linked but abolishes 5’-phosphatase activity (Mani et al., 2007). Overexpressing WT SJ1 led to an increase in LAMP2A immunofluorescence compared to the empty GFP vector expressing cells (Fig. 4A-B). In contrast, the SJ1 R258Q mutant significantly downregulated LAMP2A expression, whereas the SJ1 R839C mutant produced a less significant effect on downregulating LAMP2A (Fig. 4A-B). Interestingly, the SJ1 D769A mutation did not alter the LAMP2A expression, suggesting that SAC1 rather than 5’-phosphatase activity is involved in LAMP2A regulation. We further performed electron microscopy (EM) analysis for SJ1 R258Q-expressing cells sorted by a Fluorescence-Activated Cell Sorting (FACS) system (Fig. 4C). Compared to those expressing WT SJ1, we found the lysosome density was decreased in N2a cells transfected with SJ1 R258Q (Fig. 4D) while the lysosome size was not affected (Fig. 4E). These results, taken together, suggested that the Synj1’s SAC1 domain is important for regulating lysosome quantity.

**Figure 4.**
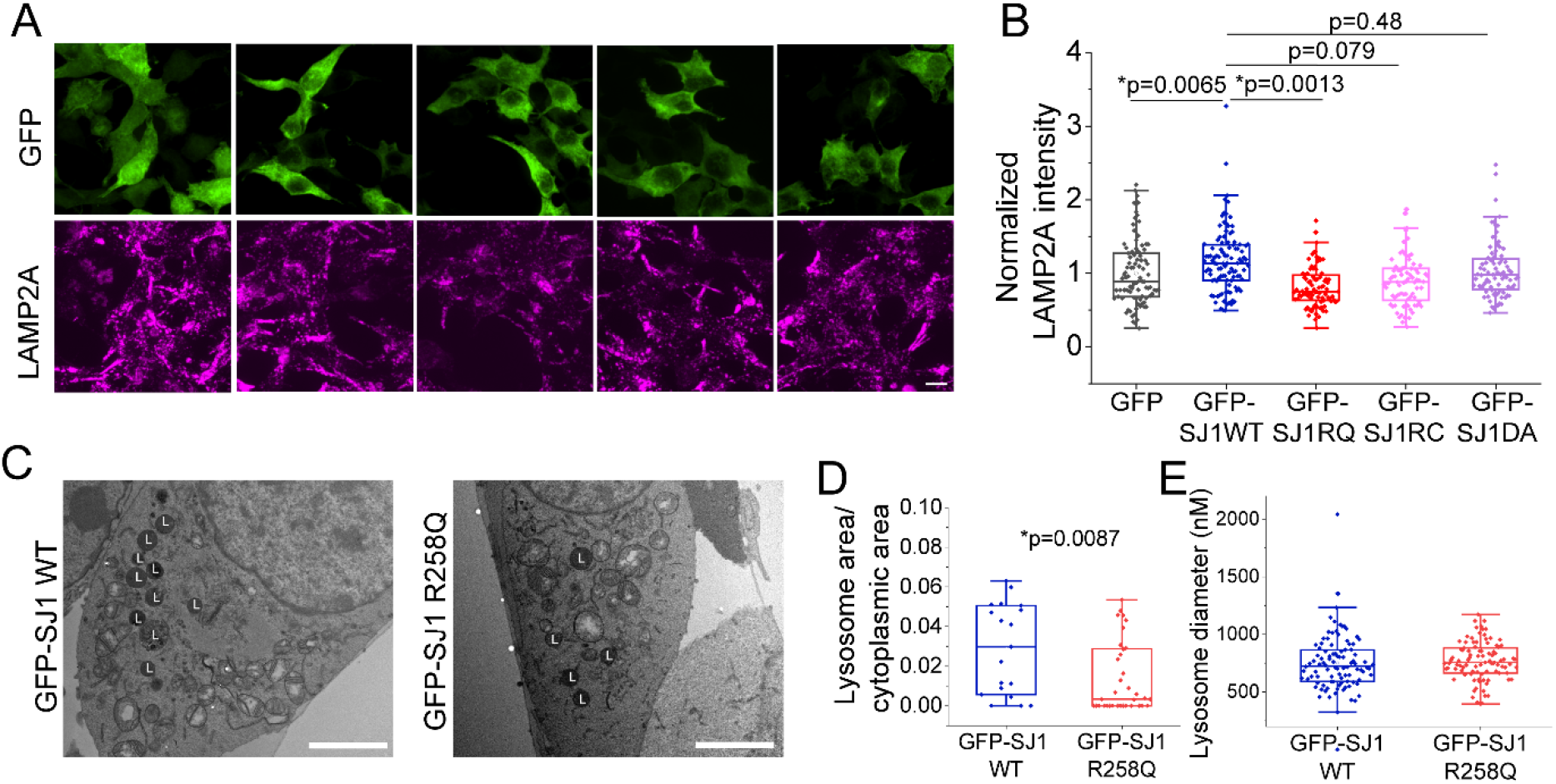
The SAC domain of Synj1 regulates lysosome abundance in N2a cells. (A)-(B) Immunofluorescence analysis of LAMP2A in N2a cells transfected with GFP, GFP-hSYNJ1 WT, GFP-hSYNJ1 R258Q, GFP-hSYNJ1 R839C and GFP-hSYNJ1 D769A. (A), representative images. (B), quantification results, N=91/104/88/80/84 (GFP/GFP-SJ1WT/GFP-SJ1RQ/GFP-SJ1RC/GFP-SJ1DA) cells. P values are from Tukey’s post hoc following One-Way ANOVA. (C)-(E) EM analysis of lysosome in N2a cells transfected with GFP-SJ1 WT and GFP-SJ1 R258Q. (C), representative EM images. L, lysosome. (D), quantification of the ratio of lysosome occupied area to cytoplasmic area in EM images for the two groups, N=19/35 (Synj1+/+/Synj1+/-) ROIs. (E), quantification of lysosome diameter in the EM images of the two groups, N=97/95 (Synj1+/+/Synj1+/-) lysosomes. The *p* value is from student’s *t*-test. Scale bar in (A), 10 μM; Scale bar in (C), 1 μM.

### Synj1 deficiency impairs α-syn A53T degradation in the axon

To determine the physiological impact of lysosomal change in Synj1+/- neurons independent of the autophagy process, we sought to investigate chaperone-mediated autophagy (CMA), which is mediated directly by LAMP2A. We hypothesized that the reduced LAMP2A would result in defective clearance of CMA substrates, such as the α-syn (Cuervo et al., 2004). We first assessed the degradation efficiency of the WT α-syn by tagging it to mEos3.2, a photo-switchable fluorescent protein. Upon UV irradiation, the green fluorescent α-syn -mEos3.2 will switch to a red-shifted protein (Zhang et al., 2012) (Fig. 5A), allowing tracking of protein degradation without interference from nascent protein synthesis. In both cell bodies and axons of Synj1+/- neurons, we did not observe significant defects in α-syn -mEos3.2 protein degradation in a 16-hour chase experiment (Fig. 5B-C), suggesting that the lysosomal defect was largely tolerated or compensated due to higher acidity. We further investigated the degradation efficiency of α-syn A53T, a prevalent mutation of α-syn found in familial PD patients (Polymeropoulos et al., 1997). We found that cell bodies of Synj1+/- neurons could clear the UV converted α-syn A53T-mEos3.2 as effectively as the Synj1+/+ neurons (Fig. 5D-E). However, the axons of Synj1+/- neurons exhibited a significant impairment (Fig. 5F-G), suggesting specific vulnerability in Synj1 deficient axons to protein stress. Taken together, our results suggest an impaired lysosomal system in Synj1 deficient midbrain neurons, which could result in a specific vulnerability in axonal protein degradation.

**Figure 5.**
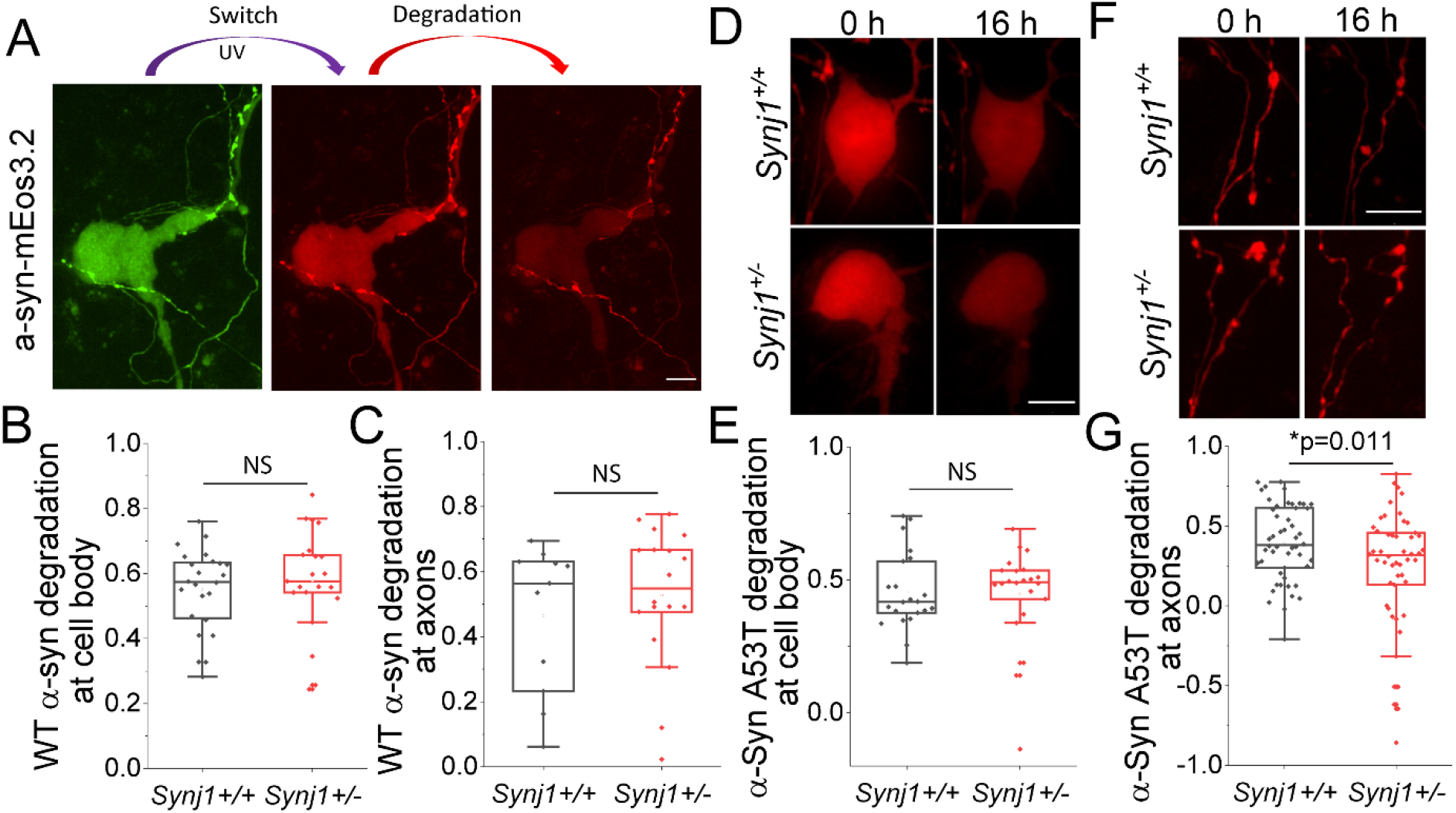
SYNJ1 deficiency impairs the degradation of α-Syn A53T in axons of cultured MB neurons. (A)-(C) The α-syn WT-mEos3.2 degradation analysis at soma and axons in cultured MB neurons. (A), representative images. (B), quantification for the fraction of degradation of α-syn WT-mEos3.2 at soma. N=24/21 (*Synj1+/+*/*Synj1+/-*) neurons. (C), quantification for the fraction of degradation of α-syn WT-mEos3.2 at axons, N=11/18 (*Synj1+/+*/*Synj1+/-*) axons. (D)-(G) The α-syn A53T-mEos3.2 degradation analysis at soma and axons in cultured MB neurons. (D), representative images of soma. (E), the degradation quantification of α-syn A53T-mEos3.2 at soma. N=22/22 (*Synj1+/+*/*Synj1+/-*) neurons. (F), representative images of axon. (G), the degradation quantification of α-syn -mEos3.2 A53T at axon, N=50/50 (*Synj1+/+*/*Synj1+/-*) axons. The p value is from student *t*-test. Scale bar in all images, 10 μM.

## Discussion

In this study, we performed a comprehensive analysis of cellular degradation systems, including autophagosomes, autolysosomes and lysosomes in cultured Synj1+/- MB neurons. We found that baseline autophagosome flux in the neuronal cell body is largely unaltered. However, we show, for the first time, that Synj1 regulates lysosomal biology. Lysosome abundance is reduced in the soma and axons of Synj1+/- MB neurons, along with a general decrease in lysosomal proteins, including LAMP1, LAMP2, and LAMP2A. We show that the SAC1 activity of Synj1 is more important relative to the 5’-phosphatase activity for regulating lysosome abundance. Interestingly, lysosomal acidity and enzymatic activity are enhanced in Synj1+/- MB neurons, which may be a direct impact of Synj1 deficiency on lysosomes or representative of compensatory change. We further examined the impact of Synj1 deficiency on cellular degradation by chasing the fluorescence decrease of exogenously expressed α-syn WT and A53T mutant proteins. We found that the degradation of WT α-syn, which most likely relies on CMA (Cuervo et al., 2004; Kuo et al., 2022) is normal in the cell body and axons of Synj1+/- MB neurons. However, the A53T α-syn exhibited impaired degradation in the axons of Synj1+/- MB neurons, suggesting an axon-specific vulnerability in removing stress proteins.

Synj1 has two phosphatase domains - the SAC1 domain enzyme that hydrolyzes PI(3)P, PI(4)P and PI(3, 5)P_2_; and the 5’-phosphatase domain enzyme that primarily hydrolyzes PI(4, 5)P_2_. Different phosphoinositides are enriched in different subcellular compartments, acting as organizers of membrane organelles and participating in cellular signaling (Posor, Jang, & Haucke, 2022). For instance, studies have shown that despite the low abundance of PI(3,5)P_2_ in cells, it is implicated in autophagosome maturation (Vanhauwaert et al., 2017) and important for lysosome formation and ion homeostasis (Ebner, Koch, & Haucke, 2019; Hasegawa, Strunk, & Weisman, 2017; McCartney, Zhang, & Weisman, 2014). Synj1 deficiency could lead to an accumulation of PI(3,5)P_2_ to interfere with lysosomal function. In our study we showed that the lysosome protein LAMP2A was decreased in Synj1+/- neurons and Synj1 R258Q-expresssing N2a cells, but not in Synj1 D769A or Synj1 R839C expressing cells. This suggests an involvement of SAC1 activity and, also, that the PI(3,5)P_2_ may negatively regulate lysosome protein expression. Indeed, PI(3,5)P_2_ is broadly involved in regulating lysosomal protein expression. It was found that PI(3,5)P_2_ can activate mTORC1 through direct binding to the mTORC1 subunit Raptor in yeast, which phosphorylates TFEB and, thereby, inhibits lysosomal gene expression (Bridges et al., 2012; Jin et al., 2014). Whether mTORC1 signaling is activated in Synj1+/- MB neurons to downregulate LAMP protein expression awaits further analysis.

Previous studies in yeast also shown that PI(3,5)P_2_ could directly bind to Vph1, a subunit of V-ATPase, to regulate V-ATPase assembly and proton pumping activity both in vitro and in vivo (Banerjee, Clapp, Tarsio, & Kane, 2019; Li et al., 2014). This is in line with our finding that lysosomal acidity is significantly lower in Synj1+/- MB neurons compared to littermate WT neurons. Interestingly, while lysosomes are much more acidic in Synj1+/- neurons than in Synj1+/+ neurons, the acidity of the autolysosomes at soma and axons measured by mKeima was not different. This could be attributed to the lack of power in the mKeima analysis, but it is also plausible that autophagosome-lysosome fusion is impaired in Synj1+/- MB neurons. For example, it was shown that PI(4,5)P_2_ not only mediates clathrin–AP2 coats assembly at the plasma membrane but also induces tubular protolysosome budding from tubular intermediates of autolysosomes (Rong et al., 2012), and participates in autophagosome and lysosome fusion (Posor et al., 2022). Whether the increased PI(4,5)P_2_ in Synj+/- MB neurons (Pan et al., 2020) plays a role in disrupting the fusion between autophagosomes and lysosomes requires further investigation.

Despite finding multiple lysosomal abnormalities in Synj1+/- MB neurons, we did not observe alteration in WT α-syn clearance, which suggests MB neurons in culture are relatively adaptive in compensating changes at the lysosomal level. The pathological α-syn accumulation in the MB of 12-month-old mice (Pan et al., 2020) may reflect age-dependent deterioration of the degradation systems or compensatory mechanisms. Interestingly, the degradation of the mutant protein, α-syn A53T, was significantly impaired in the axons of Synj1+/- neurons. Studies have shown that mutant α-syn exhibit impaired uptake by LAMP2A via the CMA pathway, which suggests evidence for a potentially defective autophagic capacity in Synj1+/- MB neurons when protein stress is present. These data are largely consistent with findings from previous in vivo studies and astrocyte cultures, which found that Synj1 haploinsufficiency minimally affects basal autophagy but neurons become more vulnerable during aging with accumulating cellular stress. We now suggest that axons of Synj1+/- neurons are more vulnerable than cell bodies, supporting the dying back theory in Synj1 deficiency associated neurodegeneration(Bernheimer, Birkmayer, Hornykiewicz, Jellinger, & Seitelberger, 1973).

## Materials and methods

### Plasmids, antibodies, and reagents

The following plasmids, pEGFP-C1-GFP-Synj1 WT, pEGFP-C1-GFP-Synj1 R258Q, pEGFP-C1-GFP-Synj1 R839C, pEGFP-C1-GFP-Synj1 D769A, pEGFP-C1-GFP-LC3 were previously reported(Pan et al., 2020; Pan et al., 2021). mKeima was gifted by Dr. Atsushi Miyawaki (Osaka University). RFP-LC3 is a gift from Dr. Huaye Zhang (Rutgers University). The plasmid expressing LAMP1-GFP is a gift from Prof. Tao Xu (Institute of biophysics, Chinese Academy of Sciences). The FIRE-pHLy plasmid was purchased from Addgene (Cat#170775,). Mouse WT or A53T α-syn cDNAs (gifted by Dr. Eliezer Masliah, National Institute on Aging) was cloned into pmEos3.2-N1 vector (from Addgene, Cat#54525) to generate the α-syn -mEos3.2/α-syn A53T-mEos3.2 plasmid.

The GlutaMAX Supplement (35050061), the Neurobasal-A Medium (12349015), the B-27 Supplement (17504044), the Basal Medium Eagle (BME) (21010046), the DMEM (11965-092), the Penicillin-Streptomycin (15140122), the Lipofectamine 2000 Reagent (11668019), the LAMP-2A Polyclonal Antibody (51-2200, 1:100 for IF), the LAMP2 Polyclonal Antibody (PA1-655, 1:200 for IF), the LAMP-1 Monoclonal Antibody (14-1071-85, 1:500 for IF), the MAP2 Monoclonal Antibody (MAP2 MA5-12826, 1:1000 for IF) and all of the Alexa Fluor dye-conjugated secondary antibodies were purchased from Thermo Scientific (Waltham, MA). The YM201636 (13576) was purchased from Cayman Chemical Company (Ann Arbor, MI). The Magic Red Cathepsin-B Assay Kit (937) was purchased from Immunochemistry (Davis, CA). The purified anti-PtdIns(3,5)P2 IgG (Z-P035, 1:100 for IF) was purchased from Echelon Biosciences. The anti-MAP2 Polyclonal Chicken Antibody (188 006, 1:1000 for IF) and the anti-MAP2 Polyclonal Guinea Pig Antibody (188 004, 1:1000 for IF) were purchased from Synaptic Systems (Goettingen, Germany). The anti-Tyrosine Hydroxylase Antibody (AB9702, 1:1000 for IF) was purchased from Millipore Sigma (Burlington, MA). The papain (LK003176) and the Earle’s Balanced Salt Solution (EBSS) (LK003188) were purchased from Worthington Biochemical Corporation (Lakewood, NJ). The Fetal Bovine Serum (FBS) (S11510H) was purchased from Novus Biologicals (Centennial, CO).

### Cell sorting and EM analysis

The N2a cells transfected with GFP-Synj1 WT or GFP-Synj1 R258Q on Day 1 were sorted to collect GFP expressing cells by an FACS equipment on day 2. Then, on day 4, the cells were fixed in the culture dish by replacing the culture medium with 2.5% glutaraldehyde in 0.1 M sodium cacodylate buffer containing 2.5 mM Ca^2+^ at room temperature. After fixation for 60-90 min, the cells were rinsed 3 times with the same buffer, and then post-fixed for 60 minutes with 1% osmium in the rinsing buffer containing 1% potassium ferricyanide. The cells were rinsed again with rinsing buffer and stained with 1% uranyl acetate at 4°C for 30 min. Dehydration with graded ethanol solutions was followed by infiltration and embedding with PolyBed812. After polymerization portions of the embedment were cut out and re-embedded in Polybed812. So, the cells could be sectioned perpendicular to the dish surface. Thin sections were viewed in a Philips CM12 electron microscope to capture images.

The diameter of lysosome was measured by drawing a line manually with the tool Straight Line in Fiji. To measure the density of lysosome, lysosome number was counted and divided by the entire cytoplasmic area defined in imageJ.

### Animal

The Synj1+/− mice were originally from the Pietro De Camilli laboratory at Yale University (Cremona et al., 1999). They were housed and handled in accordance with the National Institutes of Health guidelines approved by the Institutional Animal Care and Use Committee. C57bl/6 mice (Jax Cat # 000664) and Synj1+/- mice were used as breeders to generate Synj1+/+ and Synj1+/- pups identified by genotyping.

### Immunofluorescence (IF)

The immunostaining protocol was previously described (Pan et al., 2020; Zhu et al., 2018) with minor modifications. Briefly, the cells were fixed with 4% PFA for 15 min at room temperature (RT), permeabilized with 0.5% saponin for 15 min at RT, blocking with 5% BSA for 1 hr at RT, incubated with primary antibody overnight and incubated with fluorescent dye conjugated secondary antibody for 1 hr at RT. After complete washing with PBS, the slides were mounted with Clear-Mount (EMS, 17985-12).

### Midbrain culture preparation and transfection

The preparation of mouse ventral midbrain culture was described before (Pan et al., 2017; Pan et al., 2020). Briefly, after genotyping, the ventral midbrain tissue was dissected from postnatal day 0-1 littermates, and it was digested in a papain solution (21 U/ml, prepared with EBSS supplemented with 5 mM kynurenic acid) by stirring with a magnetic stir bar at 33-37°C for 12 min under humidified oxygenation. After spinning and discarding the digestion buffer, the collected tissue chunk was triturated by using a 1-mL pipette tip in Neuro A complete medium (0.6× Neurobasal-A Medium, 0.3× BME, 1× GlutaMAX Supplement, 1× B-27 Supplement and 10% FBS). The homogenized tissue was collected by spinning and the supernatant containing debris was discarded. The pellet was resuspended with Neuro A complete medium and subjected for spinning again. Then, the supernatant was discarded, and the pellet was resuspended with Neuro A complete medium. After counting, 25 k cells were seeded into a 6 mm × 8 mm cylinder. On day 3, medium was changed with fresh Neuro A complete medium with Ara-C (0.2 μM) to inhibit glia growth. Transfection was performed on Day 6-8 using lipofactamine 2000.

### N2a cell culture and transfection

N2a cells are cultured in 10% FBS contained DMEM medium supplemented with 10 U/ml penicillin-streptomycin. Passage procedure follows the general and standard culture protocol with digestion using 0.05% Trypsin-EDTA solution. Lipofectamine 2000 or Lipofectamine 3000 was used for plasmid transfection in N2a cells.

### Imaging and data analysis

All fluorescent images were captured using a Nikon CREST spinning disk confocal microscope with a 100× or 60× oil objective. And all the image analysis was performed by ImageJ/Fiji. For analyzing the immunofluorescence of LAMP1, LAMP2A, LAMP2 and Magic Red in neuron soma of glia-neuron culture, the floated neurons were selected for imaging. To further avoid the immunofluorescence contamination from glia, the upper confocal planes, no longer including fluorescent signal of glia, were selected for a Z Max Project performance. Then the cytoplasmic area of each neuron in the Z-MAX-Project image was circled to obtain the average value. For monitoring the fluorescent intensity of WT or A53T α-syn-mEos3.2 in neuron soma, the cytoplasmic area was circled after a Z Max Project performance to get the mean value. For ratiometric analysis of the puncta of mKeima and FIRE-pHLy as well as the IF (immunofluorescence) of LAMP1, the plugin Time series Analyzer 3.2 was used to set ROIs defined as 0.66 μM × 0.66 μM (6 pixels × 6 pixels) at the soma or axons. For analyzing the LAMP1-GFP puncta at the axons, the plugin KymoAnalyzer V1.01 (Neumann, Chassefeyre, Campbell, & Encalada, 2017) was employed to measure the puncta density. For analyzing the fluorescent intensity of a-syn/a-syn A53T at axons, the tool Segmented Line (width: 0.33 μM (3 pixels)) is used to obtain the average value through tracing the axons.

## Acknowledgements

We thank Jessica Lee, Ariana Capuano and Asma Rizvi for performing mouse breeding and genotyping to support this study. We thank Jacqueline Saenz, Hana Caiola and Mikayla Voglewede for critical reading of the manuscript.

## Figures and legends

**Extended Data Figure 2-1.**
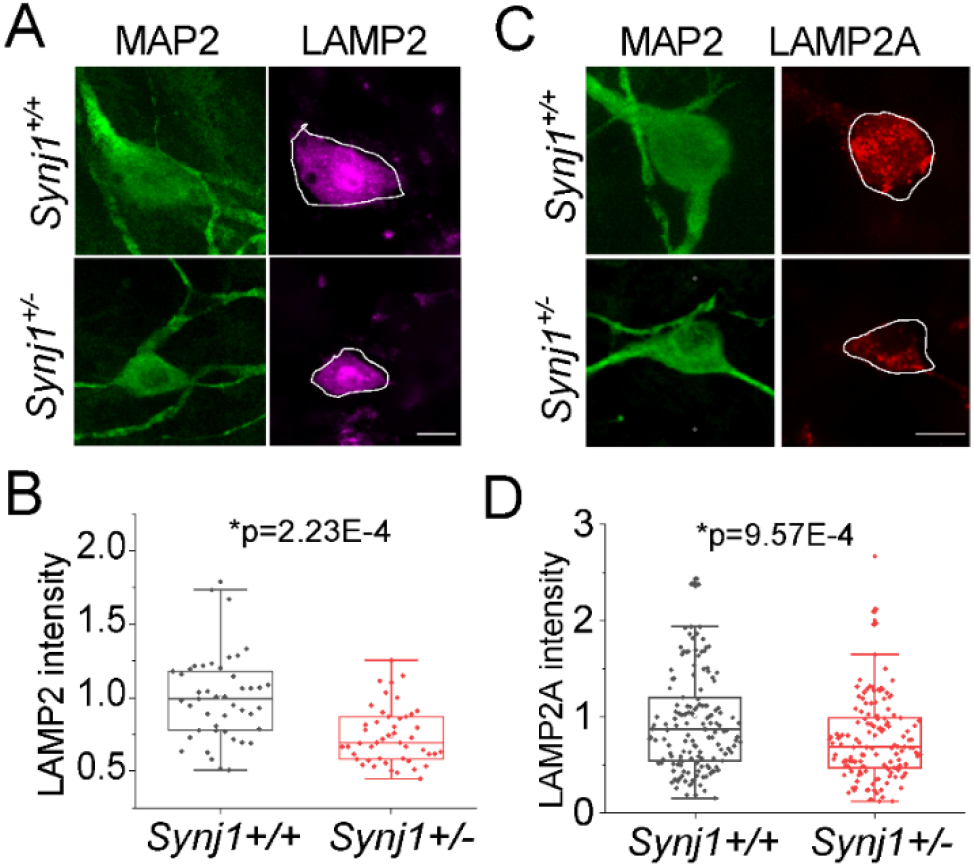
Reduced LAMP2 and LAMP2A in Synj1 deficient MB neurons. (A)-(B) Immunofluorescence analysis of LAMP2 in the soma of Synj1+/+ and Synj1+/- MB neurons, MAP2 is used as neuron marker. (A), representative images. (B), quantification results. (C)-(D) Immunofluorescence analysis of LAMP2A in the soma of Synj1+/+ and Synj1+/- MB neurons, MAP2 is used as neuron marker. (C), representative images. (D), quantification results. The *p* values for (B) and (D) are from student’s *t*-test. Scale bar in all images, 10 μM.

**Extended Data Figure 2-2.**
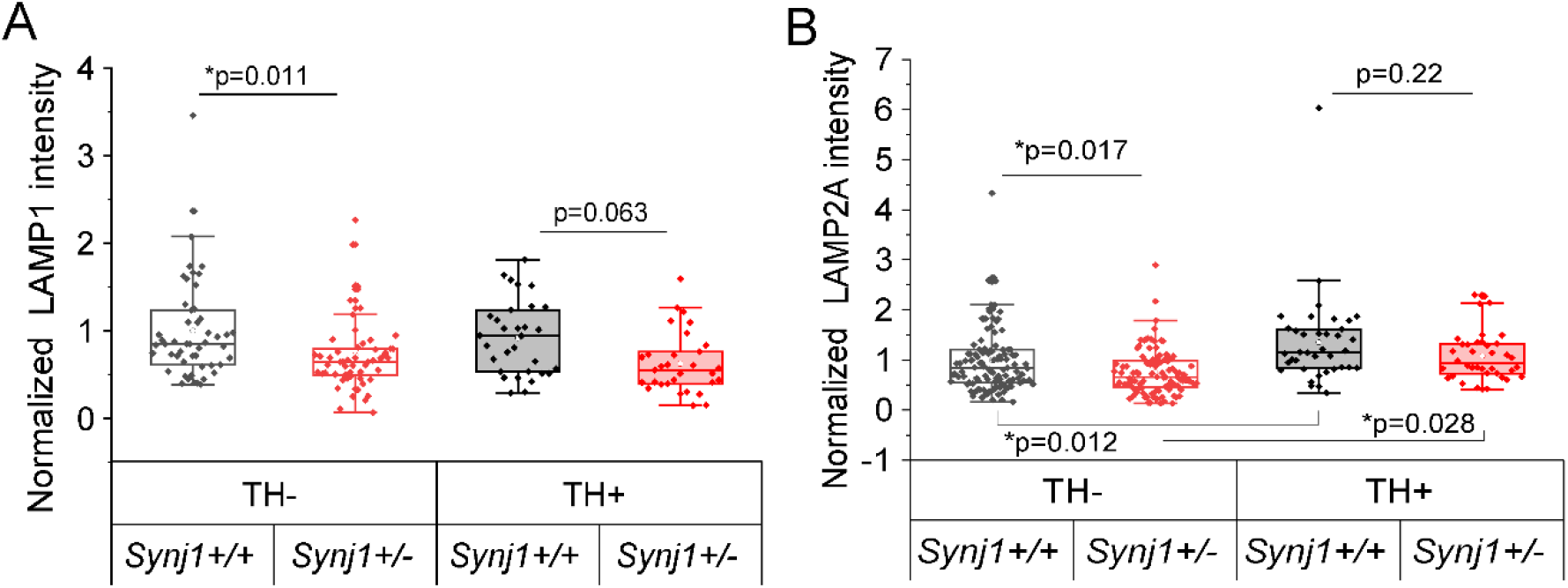
Comparison of LAMP1/LAMP2A expression between TH+ and TH-MB neurons. (A)-(B) Box plots show the expression level of LAMP1 (A) and LAMP2A (B) between TH+ and TH-MB neurons for both Synj1+/+ group and Synj1+/- group. The *p* values are from Tukey’s of Two-way ANOVA analysis.

## Notes

**Funding:** The work is supported by an NINDS R01 (NS112390) awarded to P-Y Pan.

### Competing Interest Statement

The authors have declared no competing interest.

